# Quantitative Impact of Coil Misalignment and Misplacement in Transcranial Magnetic Stimulation

**DOI:** 10.1101/2023.11.18.567677

**Authors:** Max Koehler, Thomas Kammer, Stefan M. Goetz

## Abstract

**Introduction:** Targeting in transcranial magnetic stimulation (TMS) involves the accurate placement and positioning of the stimulation coil on the head of a subject or patient. In clinical and research applications, this placement is even done manually and/or with fixed coil holders that do not compensate for motion and drift of the head. The placement involves six degrees of freedom (DOF; three position DOF: 1× contact and 2× head location; three rotational DOF: 2× alignment and 1× electric field direction/orientation), which challenge operators. This procedure is—even with an experienced user—prone to positioning errors, which can result in low treatment efficacy or high stimulation strength due to overestimating the resting motor threshold (RMT). Whereas the position and field orientation are at least widely appreciated, the coil–head alignment and its impact are often not even known. Errors involve constant errors, drift (both leading to bias and inter-individual variability), and particularly fluctuations (causing intra-individual variability).

**Objective:** We demonstrate the impact of positioning error on cortical field strength to get a better understanding of the importance of accurate positioning and compare as well as quantify the impact of position vs. alignment errors.

**Methods:** We simulated the impact in a realistic head anatomy to quantify various levels of position errors and misalignment, rolling-off the coil from the target.

**Results:** Position and alignment errors shift the focus of the electric field and reduce the electric field in the actual target. A misalignment of 10° can exceed the loss of stimulation strength in the target associated with a shift of 10 mm, corresponding to threshold stimulation leading to no detectable electromyographic response anymore. Misalignment in the direction of the handle (pitch), with which many operators appear to struggle most, reduces the field in the actual target faster than left–right roll.

**Conclusion:** This work highlights the importance of the coil–head alignment for intra- and interindividual variability.

## Introduction

Strong brief current pulses through a stimulation coil in transcranial magnetic stimulation (TMS) generate magnetic field pulses, which in turn induce electric fields and drift currents in the tissue. These electric fields can noninvasively depolarize of neurons in the brain and the periphery or muscles [1–3]. TMS is widely used in experimental brain research, medical diagnosis, and treatment [4–7]. As the most frequently used coil, the figure-of-eight coil, is rather focal, many if not all TMS procedures rely on accurate placement of the stimulation coil on the subjects_*’*_ head to ensure stimulation in the desired cortical region [8–10]. Due to the spatial organization of many functions and circuits in the brain, accurate targeting and placement is therefore essential for effective stimulation procedures and often has more impact than most other stimulation parameters [11].

Misplacement of the coil may be a key reason for low treatment efficacy [12,13]. Constant misplacement and drift may lead to bias and inter-individual variability of the outcome. Position fluctuations, which happen due to constant small head movements and neck-muscle contractions as well as due to the weight of the coil, modulate the effective pulse strength in the target from pulse to pulse, introducing intra-individual variability, which appear like an excitability change [14,15].^1^

However, coil placement errors cannot only reduce the effect of the stimulation due to insufficient activation of the required target circuits but may drive unwanted side-effects by co-activating other circuits and nociceptors [11,17]. Furthermore, placement errors can propagate to subsequent procedures. For example, the individual motor threshold estimated in a hand muscle through stimulation of the primary motor cortex is typically used for individualization of the stimulation strength also in other fields. Not considering how strongly the motor threshold may correlate or predict the necessary strength in other areas, the motor threshold is unarguably strongly dependent on coil placement. Misplacing the TMS coil during this measurement can lead to an overestimation of the stimulation strength necessary to depolarize cortical neurons, i.e., the motor threshold, and subsequent stimulation strength elsewhere [18,19]. Not only tends higher stimulation strength increase adverse effects such as scalp pain, but also increases the risk of seizure induction [20,21].

The issue of poor placement is partially known. However, the discussion almost exclusively circles around the coil position and, already rarer, the coil orientation around the head normal (yaw), which determines the direction in which the induced electric field hits a sulcus [12,18,22–30].

Placement depends on the TMS operator, experience, and most importantly awareness [18,22]. Particularly early TMS operators and but also experienced users with little work on brain targets with immediate response that give direct feedback in case of poor positioning, such as the motor cortex, tend to get the coil position and above-mentioned field orientation notably better under control than the other two rotational degrees of freedom, pitch or roll, specifically the alignment (Fig. 1). As already reported earlier, many users tend to touch the head with a point close to the handle instead of the focus area than the coil (negative pitch)[11]. As the focus area, i.e., the location on the patient-side coil surface that would generate the strongest electric field, is typically still on top of the cortical target, often thanks to stereotactical neuronavigation, the coil is only tilted towards the handle, lifting off the focus area [11]. The expected consequence is the need of a stronger stimulation strength due to the rapid decay of the field strength with distance and a lower focality [19]. In the motor cortex with direct electromyography read-out, TMS users and trainees experience the impact of the coil–head alignment rather immediate as a misalignment can almost entirely suppress any motor evoked potentials.

**Figure 1.**
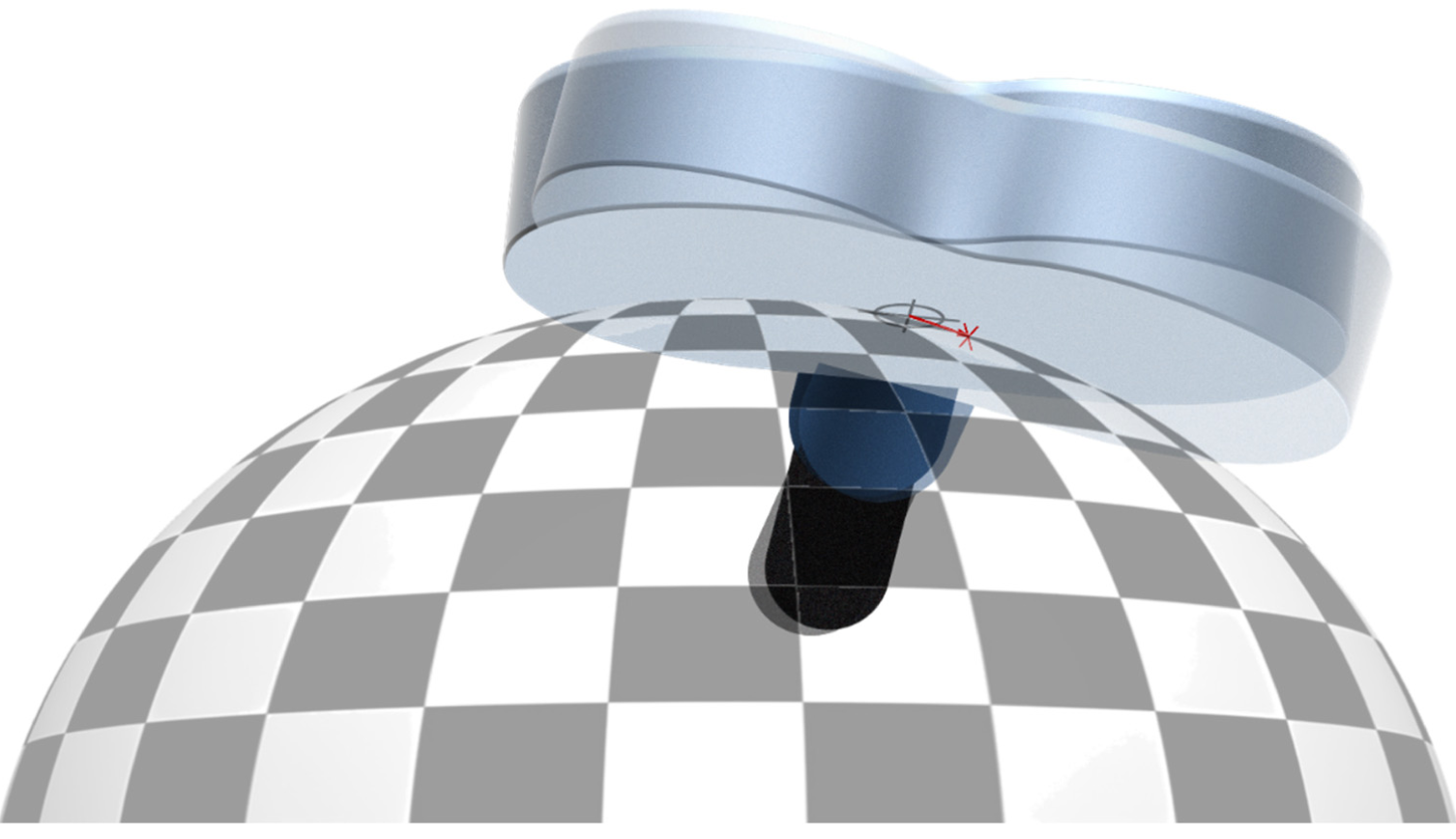
Illustration of the alignment problem when the coil is tilted. The coil rolls, i.e., does not slide, to the side so that the point of contact moves and due to the curved head surface the actual focus point, where the coil would generate its strongest electric field is lifted off.

Importantly, also robotic placement does not eliminate placement errors and often suffers particularly from bias in position and more so orientation and alignment as previous measurements demonstrate [31,32].

Despite their large impact on and variability, placement and alignment errors are not well studied yet. This analysis wants to illustrate and quantify their impact to make users aware of the problem and ideally also derive recommendations. In addition to positioning, we want to particularly include accidental roll-off of the coil from the targeted contact point on the round head surface.

## Model

We model the case of coil roll (left/right, where left is denoted as negative roll and right as positive roll) and pitch (the coil handle down, denoted as negative pitch; coil handle up, denoted as positive pitch) on the round head surface in a realistic finite-element model in SIMNIBS 4 via its scripting interface for MATLAB, using the ernie reference head model.

## Coil–head alignment

### Rotational misplacement (roll-off)

A common error pattern described earlier is the roll-off of the coil on the head surface from the focal point or desired contact point. The coil is initially placed correctly with its focal point as the contact point between the coil and the head, from which on the movement takes place.

To guarantee traceable and reproducible results and to maintain generality, we first created a parametric surface with coordinates to work on (Figure 2). The parametric surface is an ellipsoid fitted to the head marked with the 10-20 coordinate points with semi-axis lengths of a = 82 mm, b = 82 mm, and c = 106 mm.

**Figure 2.**
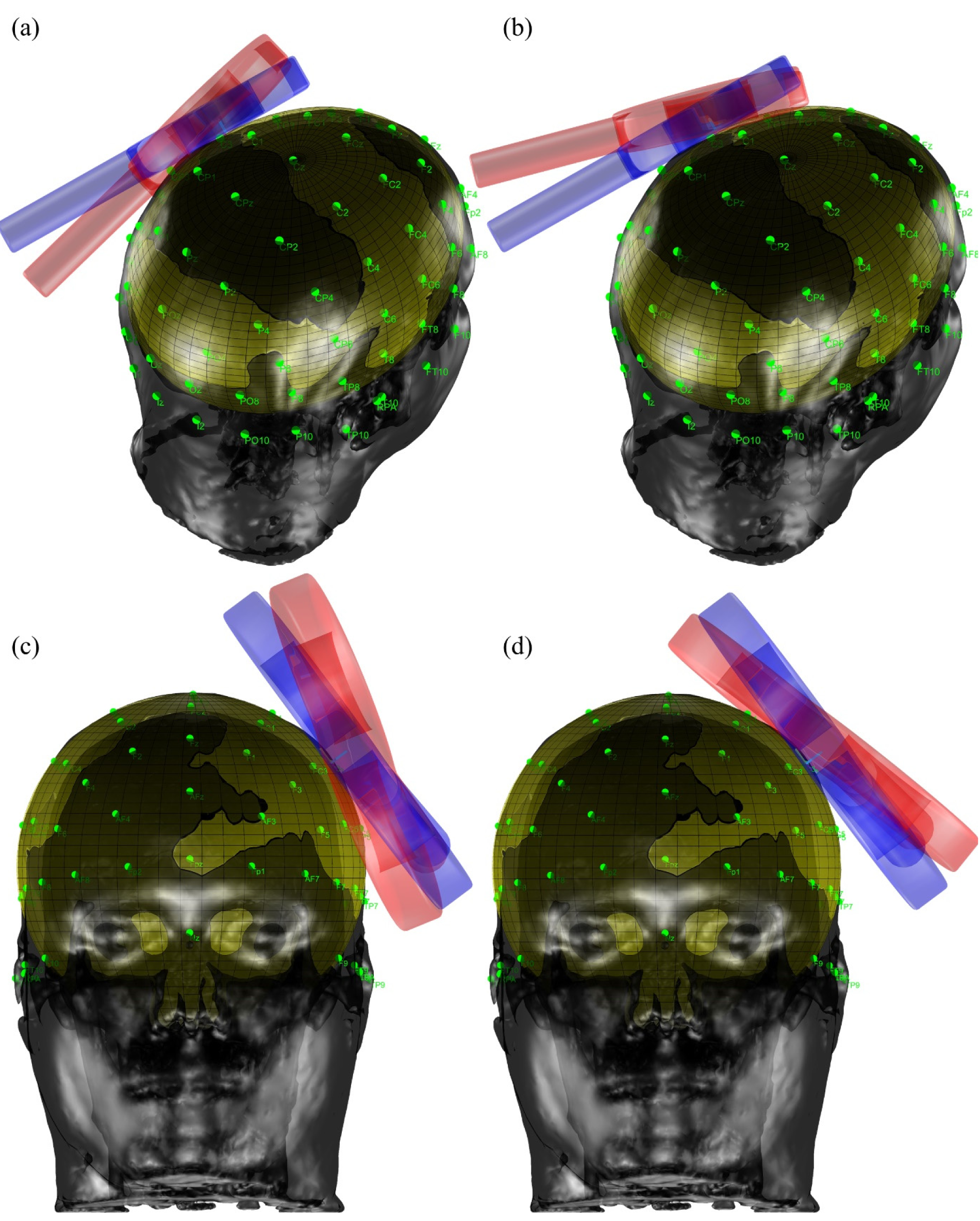
Head model with initial contact point in C3 and used ellipsoidal surface for the alignment. In (a) the coil is tilted back (negative pitch), in (b) the coil tilteed forward (positive pitch), in (c) the coil is tilted to the left (negative roll), and in (d) coil is tilted to the right (positive roll).

The normal vector of each point of the surface is the reference to for the angle of roll-off, the angle between the normal vector of the initial contact point *cp*_*n*l_ and shear of possible contact points, which are later described as a curved line. All vectors with a certain angle α revolve conically around the normal vector. This cone intersects a plane perpendicular to *cp*_l_ at cos(*α*) · *cp*_l_ with a radius of *r* = cos(*α*). Therefore, the intersecting points are

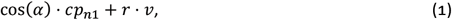

where *v* ⊥ *α* is a unit vector perpendicular to *α*. If *cp*_*n*2_ and *cp*_*n*3_ are defined in a way, so that *cp*_*n*l_, *cp*_*n*2_, *cp*_*n*3_ are orthonormal, *v* must have the form

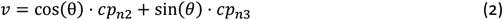

with *θ* ∈ [0,2*π*]. Later, the vectors *cp*_*n*l_, *cp*_*n*2_, and *cp*_*n*3_ form the coil coordinate system. This results in a family of normal vectors described by

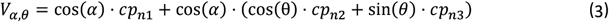

with *θ* ∈ [0,2*π*]. To transform the normal vectors back into actual coordinates, they must be multiplied with *t* = [*a*^2^, *b*^2^, *c*^2^], so that

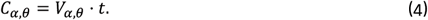

To fully determine the new contact point between the coil and the head, which we assume to be locally ellipsoidal, we also need to find the shortest distance on the surface between any point on the curve of possible points and the initial contact point, also known as a geodesic line. A geodesic is the minimization of a functional, which is the main study of calculus of variation. Having a functional of the form

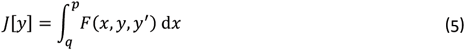

defined on a set of functions *y*(*x*) with continuous first derivatives in [*a, b*] and satisfying the boundary condition *y*(*p*) = *p, y*(*q*) = *Q*. A necessary condition for *J*[*y*] to have an extremum for a function *y*(*x*) is that it satisfies the Euler-Lagrange equation

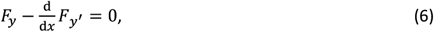

which needs to be solved. The functional we care about in this place is the arc length integral. To do so, we firstly suppose a surface **S** defined by the vector equation

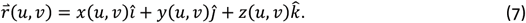

The geodesic curve lying on this surface can be specified by the equation

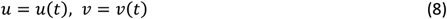

and can be obtained through minimizing the arc length integral on the near-ellipsoidal surface

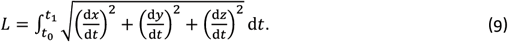

Letting 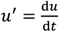 and 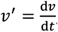, the integral can be rewritten as

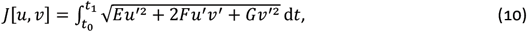

where E, F, and G are the coefficients of the first fundamental form of the surface, i.e.

The Euler-Lagrange equation corresponds to two different equations in this case:

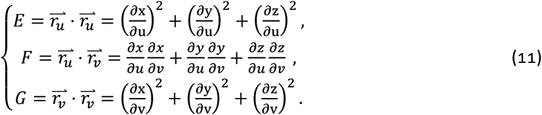

thus, we obtain

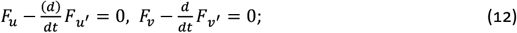

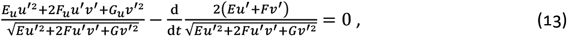

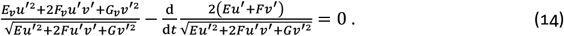

The solution of these two differential equations provides the geodesic on the surface **S**.

The following simplifications can be made in relation to our problem: *E* and *G* are explicit functions of *v* only and *F* = 0, which leads to

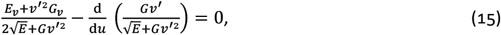

and finally to

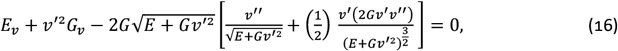

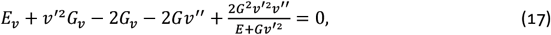

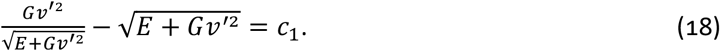

Denoting 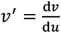 provides

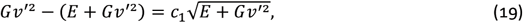

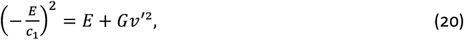

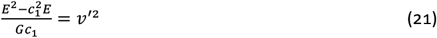

From those equations follows

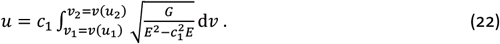

Applying this expression to an ellipsoid with the semi-axis length *a, b, c* defined by the vector equation

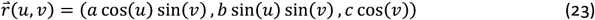

with *u* ∈ [0,2*π*) and *v* ∈ [0, *π*] resulting in the fundamental form

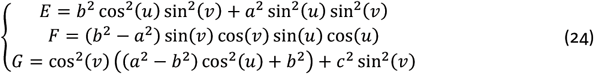

Given, that our semi-axis length a = b, the fundamental form simplifies as follows

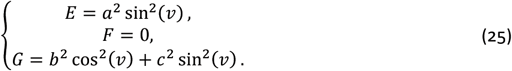

This fundamental form can be applied to the simplifications made earlier to obtain the geodesic on this surface

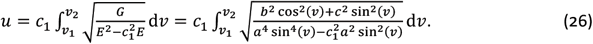

As those equations teach us the contact points and the associated arc lengths, we can determine the new position of the coil since it shifted away from the new contact point by the arc length. As mentioned before, the coordinate system of the coil is given by the normal vectors of the initial contact point, where *cp*_*n*l_ is the z-axis, *cp*_*n*2_ is the x-axis (direction of the handle of the coil) and *cp*_*n*3_ is the y-axis.

### Translational misplacement (shift)

Translational coil shifts on the surface are comparatively simple. The translation vector is added to the initial contact point in the direction of either *cp*_*n*2_ or *cp*_*n*3_ to achieve a shift in the x-or y-direction of the coil according to

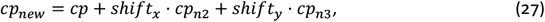

where cp is the initial contact point, *shift*_*x*_ is the shifting distance in x-direction, *shift*_*y*_ is the shifting distance in y-direction and *cp*_*new*_ the new point, where the focus point of the coil resides in 3D space.

## Coils and conditions

We chose the commercial MagVenture MC-B65 with a flat surface and the MC-B70 with angled wings as widely used figure-of-eight coils. The latter complicates the roll in the left/right direction due to a flat center strip of 34 mm width and its two wings with an inclination of 17/42.

We placed the coils on C3 in the 10–20 system with the handle pointing in the direction of P5 to represent approximately the hand representation in the primary motor cortex and pitched (up to ±30°) as well translated (up to ±30 mm) the coils in the coil coordinate system defined by the handle axis and the perpendicular surface tangent.

## Recruitment model

We incorporated a realistic recruitment model calibrated to measurements from the literature to estimate the size reduction of motor-evoked potentials [33]. The parameters representing a certain subject were (*p*_l,*i*_ – *p*_5,*i*_, *ε*_ymult,*ij*_, *ε*_xadd,*ij*_ and *ε*_yadd,*ij*_) = (–5.025, -2.235, 3.288, 1.411, 57.43, 0.07413, 5.960, 1.473·10^−6^, 0.1690).

## Hotspot detection / Hotspot determination (cortical target)

Given that the field strength in the cortical target (or hotspot) is crucial for further comparisons, we defined the target as the grey matter location below the initial contact point (C3) with the highest field strength. This location was, as expected, the most outer element of the brain closest to the initial contact point. Further comparisons are all based on the field strength in this exact target location.

## Results

As expected, both misalignment (Figs. 2 and 4) and translation (Figs. 3 and 5) lead to a reduction of the electric field strength in the target compared to perfectly tangential alignment with the coil_*’*_s focus point on top of the target. The electric field decays by approximately 10% for ∼12° forward/backward pitch and 5…10% for 10°…15° left/right roll. The decay is stronger for the angled MC-B70 as soon as the roll reached the angled wing. From that point on, the decay rate per angle almost doubles in comparison. Similar decays through coil translation require a substantial displacement of the coil: a shift of ∼10…12 mm in the handle direction leads to for approximately –5% field drop, the same shift in left/right direction to –10%. The small asymmetries stem from the change in curvature of the head towards the vertex and the asymmetry of the brain underneath.

**Figure 3.**
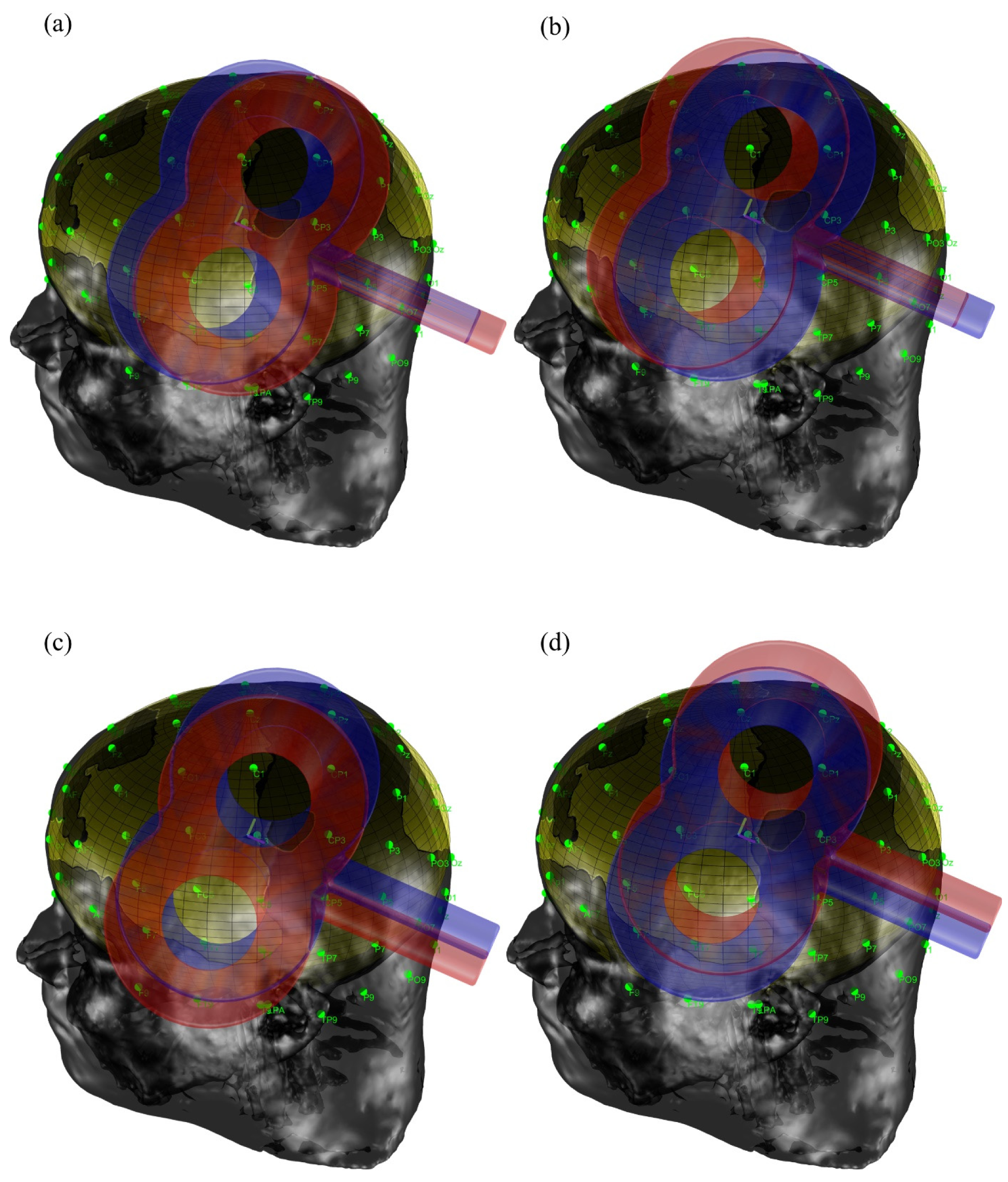
Head model with initial contact point in C3. In (a) the coil is shifted back (negative handle direction), in (b) the coil is shifted forward (positive handle direction), in (c) the coil is shifted to the left (negative side direction) and in (d) the coil shifted to the right (positive side direction).

**Figure 4.**
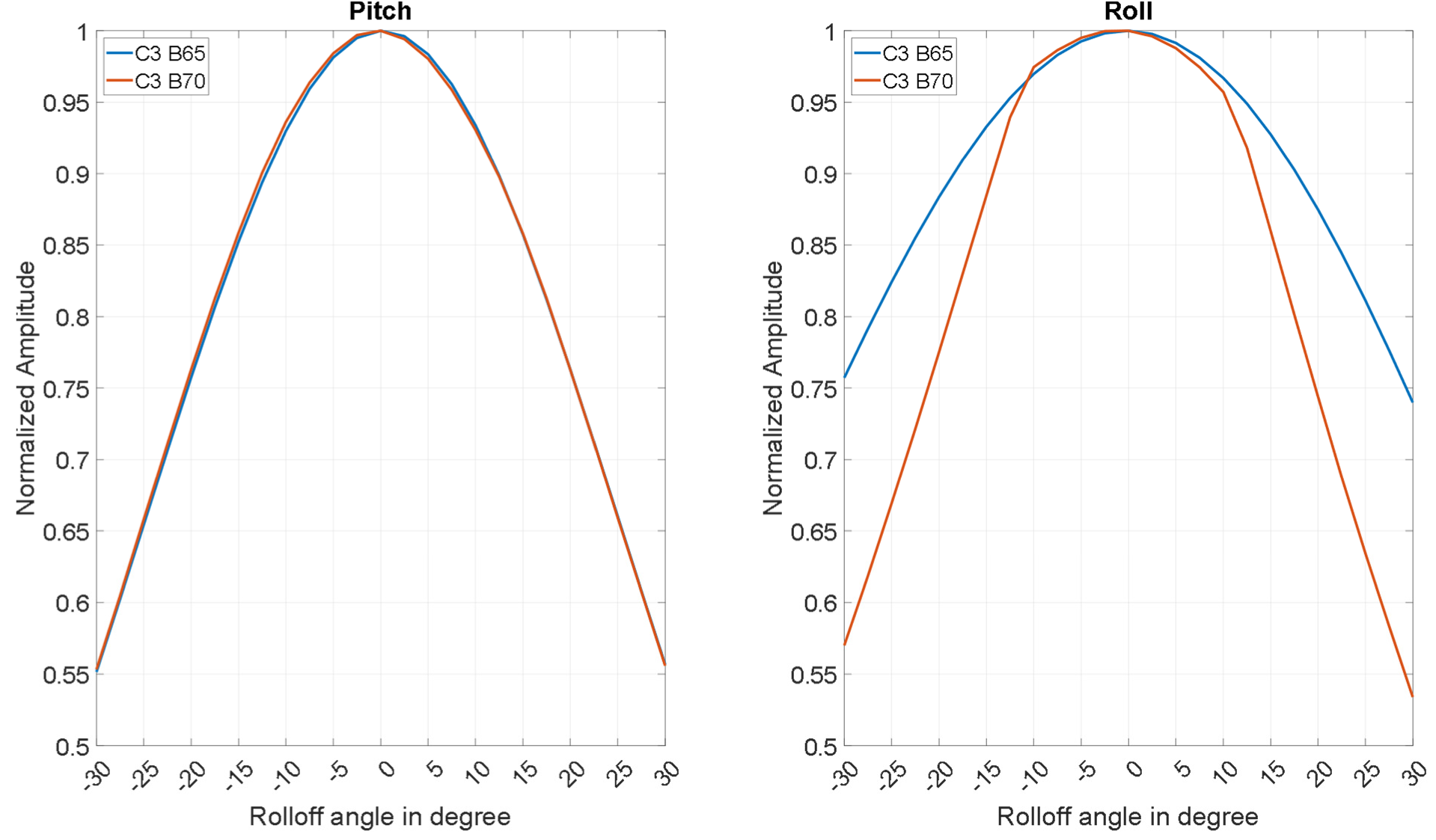
Field strength in the cortical target (hotspot) when the coil is pitched (forward/backward towards or away from the handle) or rolled (right/left) normalized to the conditions of the perfectly tangential alignment.

**Figure 5.**
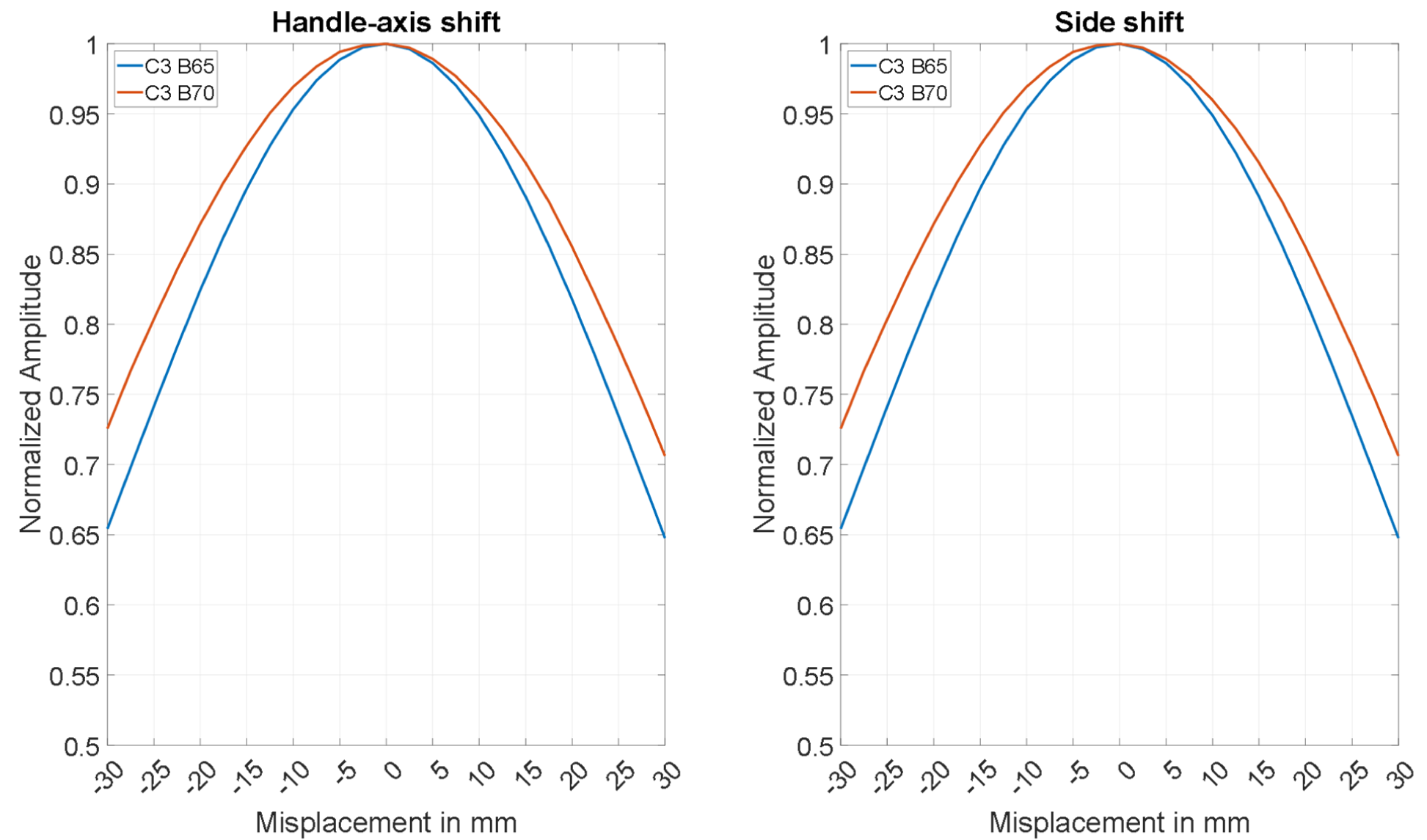
Field strength in the cortical target (hotspot) when the coil is shifted on the head surface (forward/backward towards or away from the handle or right/left) normalized to the conditions of the perfectly tangential alignment.

The steep neural recruitment translates the drop in the electric field in response to already small placement errors and fluctuations to substantial reductions in the neural output, such as motor-evoked potentials (Fig. 6). If under appropriate tangential placement on the same small hand muscle representation, the motor threshold (median of 50 μV) is reached, the evoked potentials decrease to no response (specifically the median output below the noise floor) already for less than ±10° or ±10 mm. For the 1 mV point, which is widely used for excitability probing in neuromodulation interventions, the initial distance to the noise floor may be higher, but for about ±15° or ±15 mm, the median 1 mV response is likewise vanishing below the noise floor.

**Figure 6.**
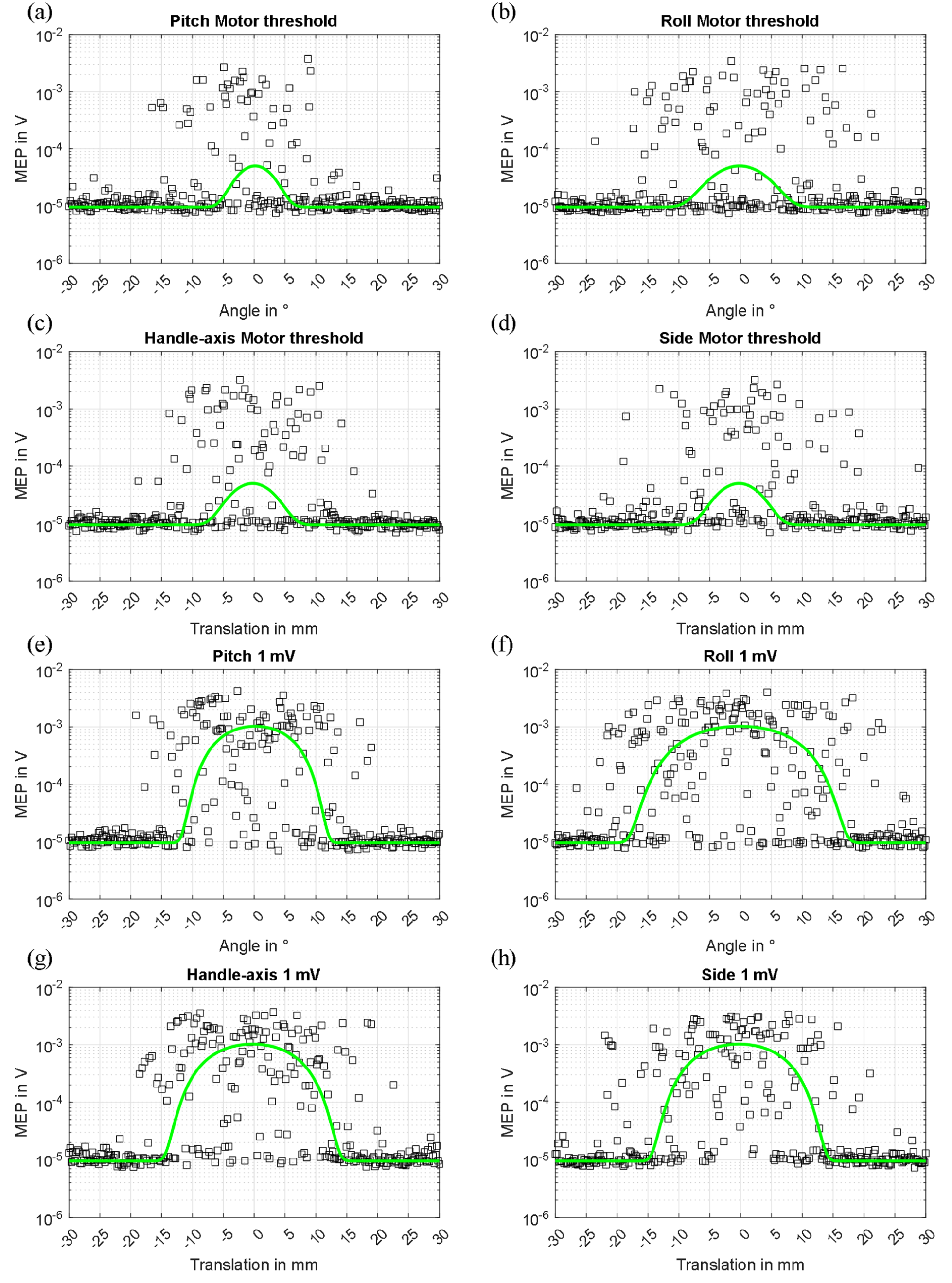
Motor evoked potentials predicted for alignment (a, b, e and f) and position (c, d, g and h) errors. The black squared indicate individual motor evoked potentials (MEP). The gree line is the median response in this position. The steep recruitment leads to a rapid drop for already small placement errors. Note the logarithmic y axis.

## Discussion and Recommendations

The paper illustrates and quantifies the field decay in the cortical target for misalignment of a flat and an angled figure-of-eight coil in a realistic head anatomy. We modelled tilting of the coil along the pitch and roll axis, as well as translation of the coil without tilt. Misalignment is among the major sources of variability for manual placement but also in case of robotic control of coil position. Dependent on the operator (or the robotic set-up and control), placement can show at least three contributions of error: (i) constant misalignment, (ii) drift, and (iii) alignment fluctuation.

(i) Constant misalignment often occurs with unexperienced users. They typically tend to touch the head with a point closer to the handle of the coil instead of a pure tangential orientation of the coil (rotation in pitch). (ii) Drifts often occur during a long-lasting session, either with coils placed manually or by use of a tripod and coil arm. Furthermore, drifts can be induced by a moving head tracker. (iii) Alignment fluctuations can occur during a session with the contact point rolling or shifting throughout an entire session.

Already moderate alignment errors have substantial impact. For the reference head model studied here, some 10° of misalignment have an impact on the same order of magnitude as a 10 mm shift. For smaller heads or different curvature, the influence of misalignment can substantially grow. These results underline that the wide spread of the field also in relatively focal figure-of-eight coils can tolerate a relatively wide shift of the coil, whereas already small misalignment on the head surface leads to larger field drops. A 10% reduction in electric field brings a threshold TMS pulse to a level where few to no motor-evoked potentials will appear in electromyography anymore.

The angled configuration of the MC-B70 coil reduces the needed stimulator power for the same stimulation strength [34]. However, the alignment issue is aggravated due to the wings compared to flat coils. On top of that, new TMS users in training often have substantially more difficulties aligning angled coils with the head as the point of contact is not as easily recognizable for the operator and can even degrade into more than one contact point when the wings touch the head.

We recommend to train new TMS users systematically in TMS coil alignment to raise their awareness. Training in the motor cortex with electromyography may help to show the strong impact or even minor roll and tilt. Several strategies can help new TMS users to test the alignment and identify where the coil touches the head: First, they can press with their thumb onto the center point of the coil on the head-averted, back side of the coil while suspending the handle and the coil cable only lightly with the other hand, which lets the coil practically self-align on the round head surface. Branding-iron coils obviously complicate this procedure. Second, a sheet of paper can be moved into the fissure between head and coil from all sides and will only move with its edge to the point of touch. If done from all sides, the point of contact can be quickly localized and visualized also during training.

Most neuronavigation systems illustrate the two rotational degrees of freedom responsible for the alignment. Typically, they are visualized as cone-representation of the normal vector or as a bubble level. However, on the one hand, most users are not aware of that feature or overwhelmed by the six degrees of freedom to control in TMS coil placement. On the other hand, this feature is only for reproducing the orientation and alignment of a previously generated target point. If the target point and alignment were generated from a previously tilted coil placement, following this guidance would lead to just a reproduction of a wrong alignment.

## Acknowledgement

The authors are inventors named on patents and patent applications for brain stimulation technology independent from this paper.

## Footnotes

1 Still noteworthy is that coil placement fluctuations are not solely responsible for pulse-to-pulse variability [16]

## Notes

### Competing Interest Statement

The authors have declared no competing interest.

